# A semi-automated technique for adenoma quantification in the *Apc^Min^* mouse using *FeatureCounter*

**DOI:** 10.1101/754325

**Authors:** Amy L. Shepherd, A. Alexander T. Smith, Kirsty A. Wakelin, Sabine Kuhn, Jianping Yang, David A. Eccles, Franca Ronchese

**Affiliations:** Malaghan Institute of Medical Research, Wellington, New Zealand

**Author notes:** To whom correspondence should be addressed at, Malaghan Institute of Medical Research, PO Box 7060, Newtown, Wellington 6242, New Zealand.

**Keywords:** ImageJ, Automated tumour analysis, Intestinal tumours, Apc^Min^, Tumour burden, Linear Discriminant Analysis (LDA), Machine Learning

## Abstract

Colorectal cancer is a major contributor to death and disease worldwide. The *Apc^Min^* mouse is a widely used model of intestinal neoplasia, as it carries a mutation also found in human colorectal cancers. However, the method most commonly used to quantify tumour burden in these mice is manual adenoma counting, which is time consuming and poorly suited to standardization across different laboratories. We describe a method to produce suitable photographs of the small intestine, process them with an ImageJ macro, *FeatureCounter,* which automatically locates image features potentially corresponding to adenomas, and a machine learning pipeline to identify and quantify them. Compared to a manual method, the specificity (or True Negative Rate, TNR) and sensitivity (or True Positive Rate, TPR) of this method in detecting adenomas are similarly high at about 80% and 87%, respectively. Importantly, total adenoma area measures derived from the automatically-called tumours were just as capable of distinguishing high-burden from low-burden mice as those established manually. Overall, our strategy is quicker, helps control experimenter bias and yields a greater wealth of information about each tumour, thus providing a convenient route to getting consistent and reliable results from a study.

## INTRODUCTION

Human colorectal cancer is a major contributor to both disease and death in the Western world, with approximately 1.36 million cases diagnosed in 2012^1^. Due to the massive impact of colorectal cancer worldwide, many animal models have been created to understand this disease and test potential treatments. Mutations in the Wingless/Int-1 (Wnt) pathway are commonplace in human colorectal cancer^2^. The Adenomatous polyposis coli (APC) protein is part of the canonical Wnt pathway, which is strongly conserved across many species, including humans and mice. APC promotes the destruction of ß-catenin and prevents Wnt signalling. Interestingly, the *Apc* gene is mutated in over 80% of colorectal cancer cases, as well as in some breast cancers^3^. One of the *Apc* mutations is particularly noteworthy, as it causes Familial Adenomatous Polyposis^4^ This hereditary genetic disease causes thousands of polyps to form in the colon of the patient, which will invariably lead to colorectal cancer if that patient is not screened and treated.

The *Apc^Min^* mouse is a widely used model of spontaneously occurring intestinal tumours that closely model human Familial Adenomatous Polyposis^5^. *Apc^M,n^* mice have been highly valuable in demonstrating key mechanisms in colorectal cancer, for example, the importance of Vascular Endothelial Growth Factor in the initial growth of intestinal tumours^6^, the role of COX-2 in adenoma formation^7^, and the role of IL-33 in promoting tumorigenesis by modifying the tumor immune environment^8^. *Apc^M,n^* mice produce an inactive, truncated APC protein due to a mutation leading to a premature stop codon in the *Apc* gene^9^. This functional loss in *Apc^Min^* mice favours aberrant cell growth and, ultimately, spontaneous adenoma generation in the mouse intestinal tract. Adenomas continue to grow throughout the mouse’s life, eventually causing bleeding, anaemia, and death, suggesting that tumour size, rather than tumour count, may be a relevant metric.

Despite the wide use of the *Apc^Min^* model, there is no standardized technique to quantify adenoma burden in these mice. Most papers rely on complex protocols and report only on manually-counted adenoma numbers, or numbers and areas in selected areas of the intestinal tract, although some also include information on adenoma location and size. However, high quality semi-automated methods are now becoming available to facilitate the identification of tumour lesions in histological images^10^, or guide the visual classification of macroscopic tumour lesions including melanomas in patients^11^. Therefore, these methods can offer rapid and objective tumour identification in a broad range of situations.

In this paper, we describe a protocol for preparing standardised, photography-based images of mouse small intestine (SI), large intestine (LI) and caecum; a new ImageJ^12^ software macro called *FeatureCounter* that automatically identifies tumour-like features in the SI images and extracts measures such as area; and a machine learning pipeline for classifying these features as true adenomas or not. We illustrate this strategy’s performance on 120 mice of different genotypes, age and sex. On the whole, our approach extracts a more detailed picture of the adenoma burden in mice in a standardized and reliable manner, enabling a rapid and more sophisticated analysis of the experimental results.

## RESULTS

### Adenoma enumeration approaches

Unbiased and reliable evaluation of tumour burden is essential to the interpretation of the results of any preclinical study addressing tumour biology and potential therapy. This is normally achieved by blinding investigators to the treatment group and performing lengthy manual quantification under a microscope. Nonetheless, individual variations in measurement techniques make the standardization of results across different investigators difficult to achieve. To overcome these limitations, we designed three new techniques to evaluate tumour count and area in the SI of tumour-prone *Apc^Min^* mice. The three techniques differed in degree of automation, in how “features” of interest were identified, and in how those features were classified or “called” as true Adenomas or not. A diagrammatic representation of the steps and approximate time taken to perform a traditional method and these three new techniques are shown in **Fig. 1**. A summary of these approaches is provided below:

1. The TRAD (Traditional) method involved dissecting the intestinal tract, longitudinally opening the gut, spreading the tissue onto a petri dish or glass plate, and manually enumerating tumours on fresh tissue using a stereomicroscope (**Supplementary Fig. 1**). The nature of these visually-identified tumours can be confirmed by standard histological techniques as shown in **Supplementary Fig. 1**.
2. The DRAW approach involved dissecting the SI and removing all fat tissue, opening it longitudinally taking care to leave any visible tumours intact, and carefully spreading the tissue flat on a suitable cardboard as detailed in the Methods section. This was then photographed close-up with a white ruler in shot for scale, and the photo stitched together and opened using the Java-based image processing programme ImageJ^12^. The image was scaled using the ruler, and the ImageJ ‘freehand selection’ function was used to manually draw the margin of each of the visually-identified Ad. These features were then measured and added up using the ImageJ’s ‘analyze particles’ function to generate adenoma numbers and area. The same approach was used also to quantify tumours in the LI and caecum, after they were prepared similarly to the SI. In this study, the DRAW approach identified no adenomas in the SI, caecum or LI from control mice, indicating high researcher reliability when identifying tumours.
3. The CALL approach followed the DRAW approach up to the full SI image opening in ImageJ. At this point, the *FeatureCounter* macro was run in ImageJ to automatically set the scale and outline the contour of interesting features that might be adenomas. From here, a researcher manually located each feature and “<u>called</u>” (assigned) them as ‘true Adenomas’ (Ad) or ‘not Adenomas’ (nAd). The resulting information is used by ImageJ ‘analyse particles’ function to calculate adenoma number and areas. Thus, the CALL approach automatically identifies adenoma-like features that are then verified by eye, providing a gold-standard training set for machine learning if required.
4. Finally, the LDA approach used the *FeatureCounter* macro-identified features generated using the CALL approach and <u>L</u>inear <u>D</u>iscriminant <u>A</u>nalysis (LDA, a simple machine learning technique) to determine how to discriminate between Ad and nAd features based on the feature measures. Once trained on a CALL dataset, this method is fully automatic, and features can be delineated by *FeatureCounter* and then classified as an Ad or nAd by the LDA.

**Figure 1.**
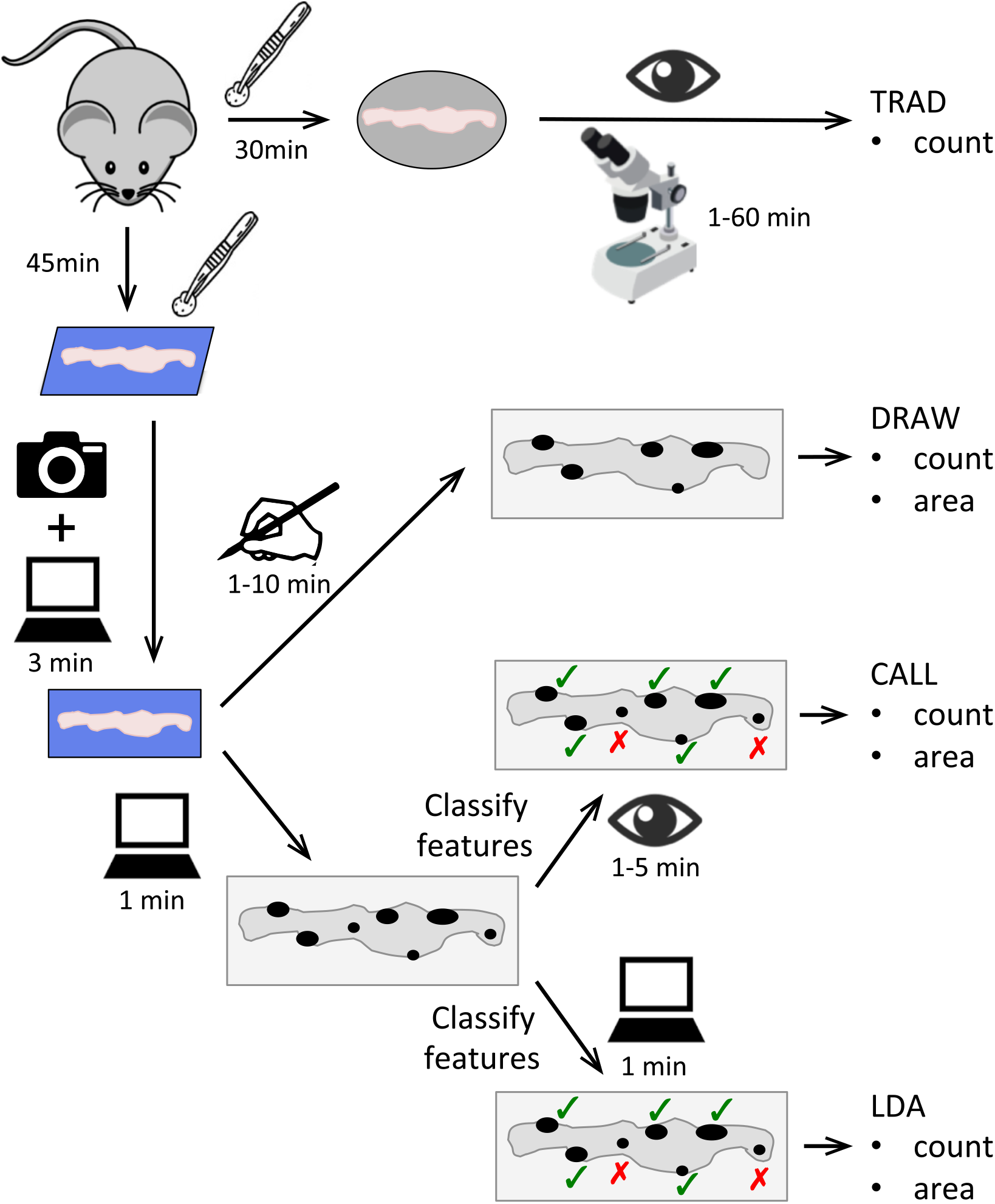
Schematic of the tumour measurement methods described in this paper. Flow chart illustrating each step needed to perform the TRAD, DRAW, CALL, and LDA intestinal adenoma identification methods described in this paper. The icons represent the tools required to perform each step; estimated time costs per step are indicated. Please refer to the Materials & Methods and Results for a detailed description of the workflow for each method.

### Photography and *FeatureCounter* can be faster than manual quantification

We compared the time required to quantify SI tumours using the various approaches described in **Fig. 1**. Preparing the SI for analysis using the TRAD approach took about 30 minutes. In contrast, the time to prepare and photograph one SI sample for all other approaches took in total about 40 minutes, including sample dissection, washing, photographing, and image stitching time. Similar quality images were obtained using either fresh SI tissue or tissue that had been stored frozen and thawed before sample processing and analysis. Use of frozen tissues added about 5-10 minutes to the total tissue preparation time, but introduced a very useful experimental breakpoint option when immediate analysis was not possible or highly inconvenient, as is often the case in survival studies.

The quantification of tumours using the TRAD approach, by visually quantifying tumours under the dissecting microscope, took up to 60 minutes per sample depending on tumour burden. Measurement of individual tumour sizes would add considerably to this time, especially when the tumour burden is high. In the DRAW approach, tracing features by hand in ImageJ took about 1 to 10 minutes per sample, again depending on tumour burden. Running the *FeatureCounter* macro to automatically identify image features of interest took about 15-30 seconds. Manually calling tumour features from the *FeatureCounter* macro’s features in the CALL approach took 1 to 5 minutes per sample, while the LDA approach (assuming a streamlined processing pipeline) took only one minute to complete the analysis across all 3188 features from 117 animals. It is immediately apparent that the main time gain is in the ability to automatically identify and call features, which is highest on heavily tumour-burdened mice. For low-burden mice, the extra preparation time would offset this gain; however, the consistency and depth of data generated using the DRAW, CALL or LDA methods may make the extra time investment beneficial compared to the TRAD approach. Overall, the TRAD approach takes approximately 90 minutes per sample, the DRAW approach 60 minutes, the CALL approach 50 minutes and the LDA method 45 minutes per sample. **Figure 1** schematizes these four approaches along with time costs for each step of each method.

### Tissue preparation and *FeatureCounter* True Positive Rate

High quality tissue preparation is essential to tumour identification using the *FeatureCounter* macro. **Figure 2A** shows a SI laid out on cardboard, before being bisected into two long pieces which were then cut longitudinally and, using tweezers, opened out, spread flat with smoothed edges, and cleaned with PBS to expose any adenomas present. A representative image is presented in **Fig. 2B**. Tumours are visible as denser white areas on the blue cardboard background. From these images, tumours were manually delineated by an experienced researcher to generate the DRAW mask in **Fig. 2C**. Alternatively, the *FeatureCounter* macro was used to automatically flag adenoma-like areas and generate a mask as shown in **Fig. 2D**. *FeatureCounter* identified very few features from a good preparation of control SI with no adenomas. Representative image and mask are shown in **Fig. 2E** and **2F**, respectively. Common issues with tissue preparation and image analysis include rolled edges, excess fat, patches of dried tissue, and light reflections which can all be picked up as non-tumour features by the *FeatureCounter* macro (**Supplementary Fig. 2**). These “false positive” image features can be largely avoided by first removing excess fat at sample collection and then, during preparation, ensuring that the tissue edges are flat by smoothing with tweezers, regularly moistening the samples once mounted, and finally ensuring consistent camera and light placement during photography. Once the protocol is learnt, it is relatively simple to avoid all these artifacts.

**Figure 2.**
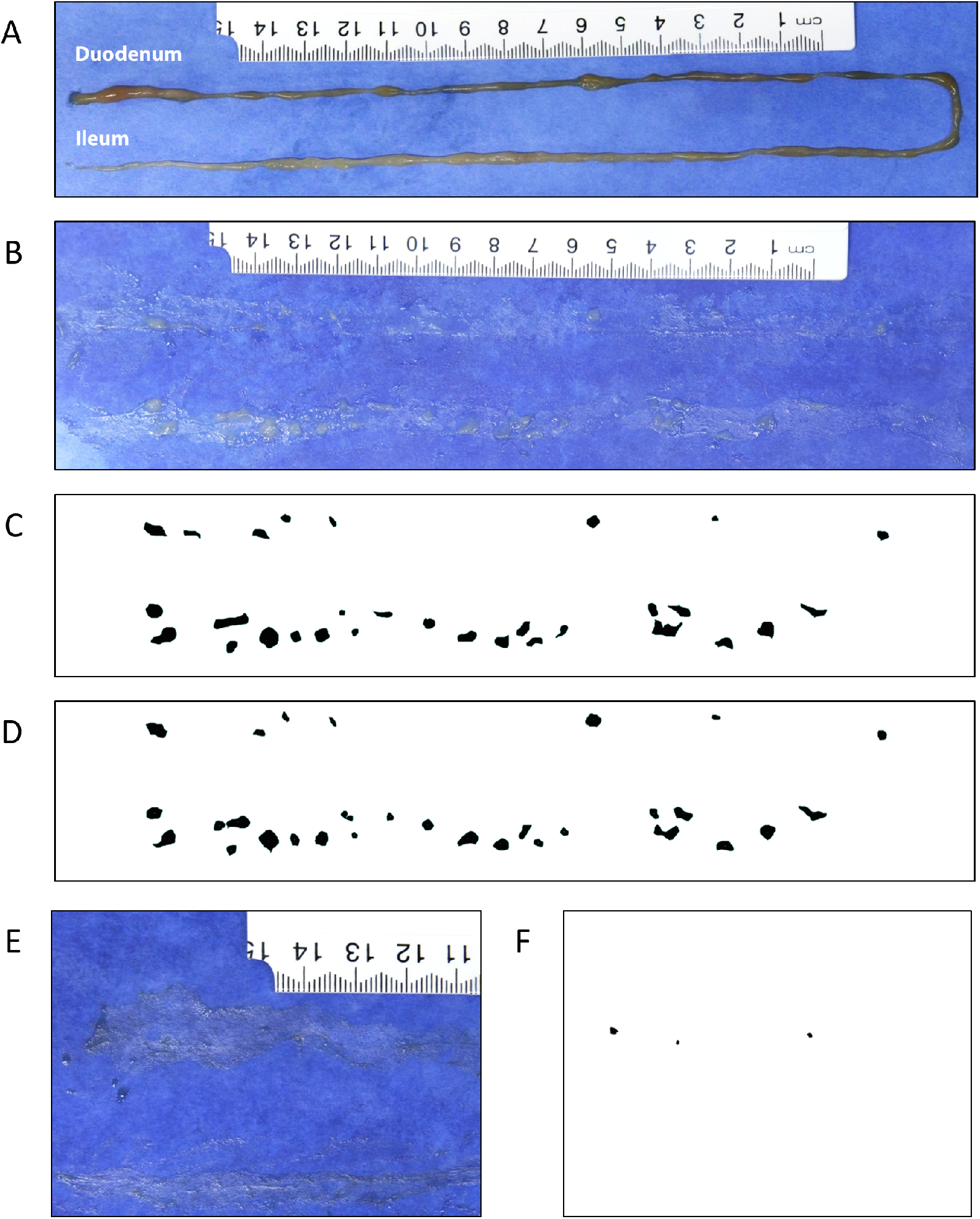
The image features (adenomas) identified by the automated *FeatureCounter* macro mostly correspond to adenomas as identified using the manual DRAW method. (**A**) Freshly collected SI from an *Apc^Min^* mouse placed on blue cardboard. (**B**) The same SI after being cut longitudinally, spread and and cleaned with PBS to expose tumours. (**C**) *FeatureCounter-generated* tumour mask for the same sample. (**D**) Manually-generated tumour mask for the same sample. (**E**) A representative partial picture of a control SI. (**F**) *FeatureCounter-generated* mask, showing features picked up on the section shown in E. No additional features were picked up from the complete image.

### Validation of tumour identification in the small intestine

To ensure that our premise of identifying image features as actual adenomas was correct, we carried out experiments where fresh SI tissue was spread on blue cardboard, analysed using the DRAW method, and then used as a source of tissue for microscopic analysis. As shown in **Fig. 3C** and **3D**, two putative adenomas were selected due to their relatively isolated location away from other tumours in the same sample, removed using a scalpel, then formalin fixed, embedded in paraffin, and stained with haematoxylin and eosin. **Figure 3A and 3E** show a magnification of these adenomas. Microscopic images in **Fig. 3B and 3F** revealed a typical morphology with thickened mucosa, glandular appearance and a sessile structure. This appearance is characteristic of adenomas as described in *Apc^Min^* mice^5^ and very similar to that of *Apc^Min^* adenomas imaged in our Lab using standard methods such as Swiss rolling of intestinal tissue (**Supplementary Fig. 1**).

**Figure 3.**
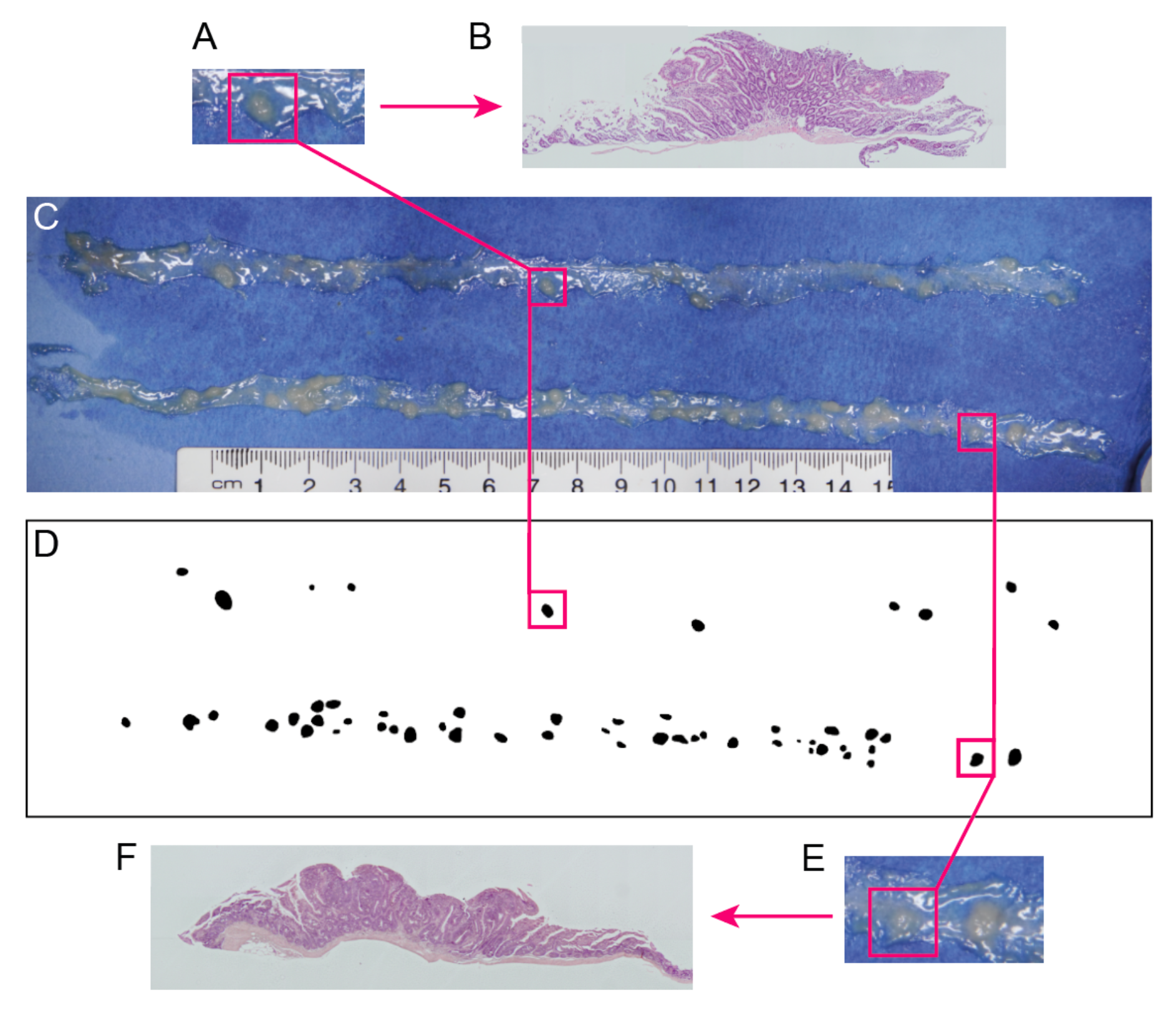
The image features identified using the DRAW method are adenomas. Fresh SI tissue was isolated from *Apc^Min^* mice, immediately set up on blue paper (**C**) and examined using the DRAW method in *FeatureCounter* to generate the mask in (**D**). Two relatively isolated features were chosen (marked by orange lines and magnified in **B, E**) excised from the paper support using a scalpel, and processed by formalin fixation, paraffin embedding and H&E staining to generate the images in (**A**) and (**F**). Data are from one of 3 mice and 7 SI tumours that were similarly treated and analysed.

As a further validation of the tumour-bearing status of *Apc^Min^* mice as determined using the DRAW method, we compared spleen and body weight between groups of *Apc^Min^* mice and their adenoma-free WT littermates, which were sacrificed at the same time or shortly after euthanasia of the last surviving *Apc^Min^* mouse in the same litter. A total of 49 mice, 27 *Apc^Min^* and 22 WT, were assessed. The average age of the *Apc^Min^* mice was 149 days with SD of 37, while the average age of the WT controls was 177 ± 21 days. The results in **Fig. 4** show that spleen weight was significantly higher in *Apc^Min^* mice compared to WT controls, while body weight was lower. This is consistent with the reported anemia that develops in *Apc^Min^* mice with increasing tumour burden, which in turn leads to splenomegaly^5^. All *Apc^Min^* mice harboured numerous adenomas in the SI and a considerable tumour burden measured as total tumour surface throughout the SI. No tumours were detected in the WT littermates.

**Figure 4.**
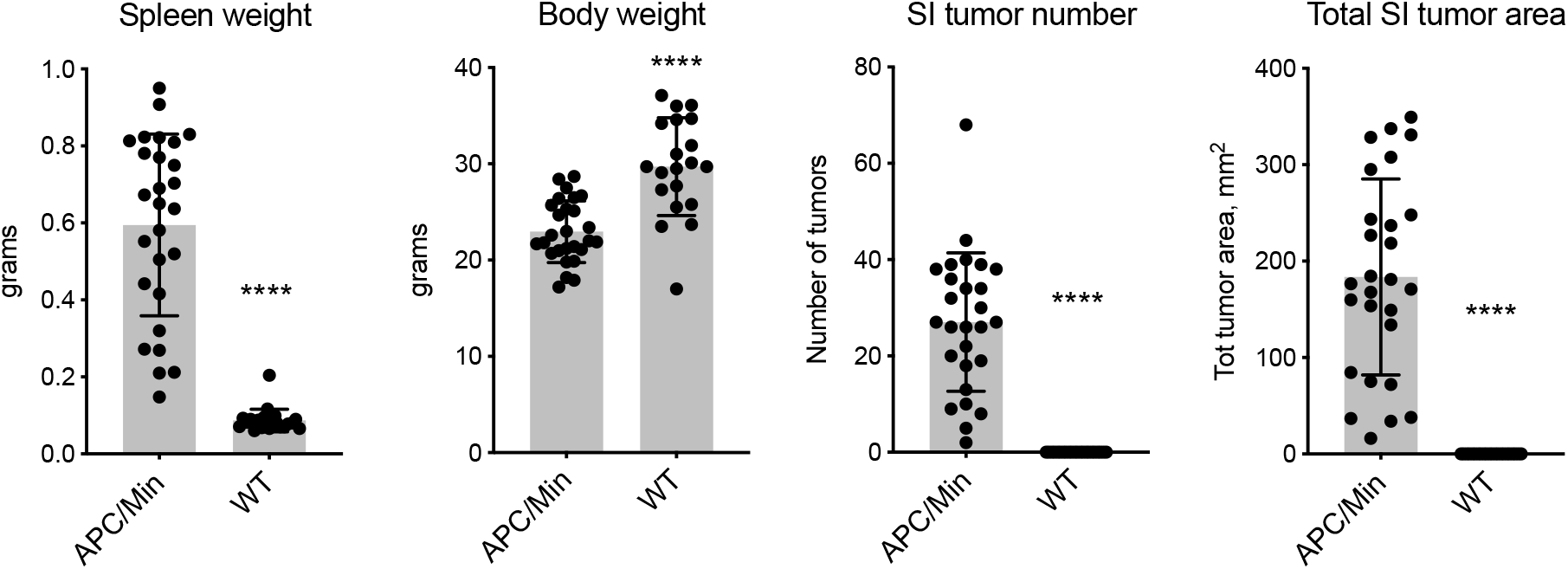
Spleen weight, body weight, number of SI tumors and their total area differ significantly between *Apc^Min^* mice and their WT littermates. *Apc^Min^* mice (n=27, 13 females and 14 males) were sacrificed when anemic and their body and spleen weights were determined. SI tumor numbers and total area were determined as shown in Figure 2 using the DRAW method. WT littermates (n=22, 9 females and 13 males) were sacrificed together with, or soon after, the last surviving *Apc^Min^* littermate. Average ages ± SD were 149 ± 36 days for *Apc^Min^* mice, and 177 ± 21 days for WT controls. Bar graphs show mean ± SD, each dot represents one mouse. P values were calculated using a Mann-Whitney test, ****: p<0.0001.

### Linear Discriminant Analysis setup and feasibility

We postulated that it would be possible to identify the true adenomas amongst the SI image features delineated by *FeatureCounter* using data from the 22 shape and colour feature measures provided by ImageJ. For example, one might expect adenomas to have rounder shapes and slightly different colour than fat deposits and other non-tumour features. We thus investigated the use of machine learning techniques for separating the true adenomas, “Ad”, from not true adenomas, “nAd”. To provide a full training data for such a classifier, all the image features from 120 mice with complete measures were called as Ad or nAd by a blinded, experienced researcher using the CALL method. The dataset ultimately contained 3447image features (1286 Ad, 1919 nAd, rest unclassified).

As a first analysis, we performed a PCA of the of the image feature data generated using *FeatureCounter.* It was quickly apparent that there was segregation – though imperfect – between the Ad and nAd classes (**see Supplementary Fig. 3**), suggesting that it was likely that the LDA would be able to identify true Ad from nAd. We thus pursued the LDA to try and automatically separate the feature classes based on the measure data.

Non-independence of observations can be a major problem in any statistical methodology not designed to take it into account, as is the case for LDA. Here, observations (image features) are nested within mice, in other words, many features may be found in the same mouse, potentially causing non-independence of observations. This may be an issue if, for example, a generally low-quality gut preparation led to bias in one or more image feature measurements across all features from that mouse: the LDA learning would include this bias and thus fail to generalize properly to all features. We thus used the PCA in **Supplementary Fig. 3** to highlight potential mouse-level biases. As shown in **Supplementary Fig. 3**, the barycentres of most of the 120 mice clustered at the center of the PCA, indicating no major mouse-level bias. For animals with barycentres not clustering within this central area, SI photographs were retrieved and scrutinized for signs of substandard preparation. We concluded that 3 mice had photography of insufficient quality due to either poor sample preparation or inappropriate camera settings. After excluding these, no such bias was observed. This result emphasises the importance of standardising the tissue preparation and photography protocols to minimise sample batch effects. After this step, 3188 features with proper CALL classifications (1279 Ad (40.1%) and 1909 nAd (59.9%)) from 117 mice were retained for training the classifier.

### Linear Discriminant Analysis performance

As with any classification strategy, it is good practice to perform a validation experiment to assess the classifier’s stability and performance when faced with novel data; in other words, we wanted to check that the LDA classification strategy would perform well when applied to real-world experimental numbers. Using a “bootstrapping” random sampling with replacement strategy (see LDA validation in Methods), we generated a total of 4000 validation datasets, computationally representing the equivalent number of ‘experiments’ of normal *Apc^Min^* and WT animals, and each was used to train a separate LDA. We chose a bootstrapping approach due to the relatively smaller size of our dataset, and selected with replacement to ensure that population distribution was maintained for selections within each validation dataset. For each validation set, feature-level performance indicators including accuracy, TPR and Positive Predictive Value (PPV, or precision) and dataset-wide performance indicators (such as the ratio of positive adenoma calls over true adenomas) were derived for Ad and nAd on the full dataset, and compared to those obtained using LDA on the full dataset, as described in the Methods.

The distributions of the feature-level performance indicators are presented in **Fig. 5A and 5B**. The accuracy achieved for the full dataset was of 87%. The TPR (or the percent of the true Ad / nAd correctly identified as such by the LDA) for the full LDA of Ad and nAd were close to 80% and 90%, respectively, indicating that the LDA was identifying correctly the majority of both real Ad and nAd features. The PPV (or the proportion of the features identified as Ad / nAd by the LDA that were correctly identified) of Ad or nAd were approximately 85% and 87% respectively, again showing good performance of the full LDA to classify Ad and nAd features. Unsurprisingly, the LDA done on the whole dataset outperformed the majority of bootstrapping datasets, perhaps indicating a slight overfitting when using the full dataset. Nonetheless, all the indicators obtained on the validated datasets remained strong (indeed, the worst performing indicator was Ad.TPR, with only 75% of values above 75%).

**Figure 5.**
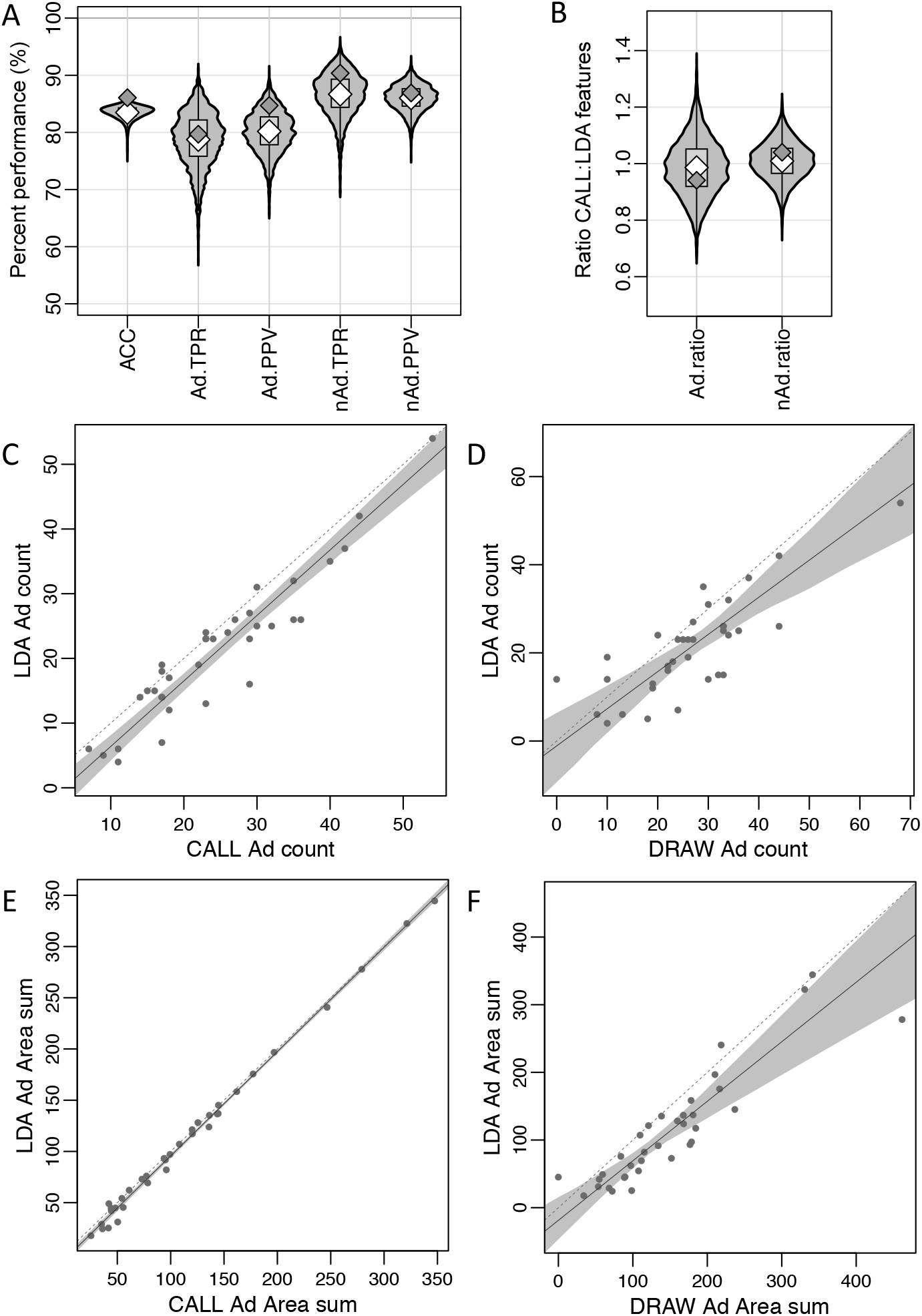
LDA predicts *Apc^Min^* tumour count and area with similar accuracy to the CALL and DRAW approaches. (**A, B**) Violin plots illustrating the distribution of the selected LDA performance indicators across the 4000 cross-validation datasets from 117 mice, each including 750-959 image features, when compared to the CALL-defined adenomas. The light grey violins are representative of the distribution of values obtained across the CV datasets; central grey boxes indicate the middle 50% of values; white diamonds represent median values for the CV datasets; dark grey diamonds represent the values observed in the full LDA. (**A**) shows Accuracy (ACC); Ad True Positive Rate (TPR, or sensitivity); Ad Positive Predictive Value (PPV); nAd TPR (or specificity); and nAd PPV distributions. **(B**) The Ad.ratio is the ratio between the number of CALL Ad and LDA Ad, with a value of 1 indicating a perfect match. The nAd.ratio is determined similarly for nAd features. (**C-F**) Deming regression plots comparing mouse-level adenoma number and total area values obtained through different approaches, for 35 mice. (**C**) compares adenoma counts generated by the LDA and CALL methods, (**D**) compares adenoma counts generated by the LDA and DRAW methods, (**E**) compares total adenoma area generated by the LDA and CALL methods, and (**F**) compares total adenoma area generated by the LDA and DRAW methods. Each dot corresponds to one mouse. Dotted grey line represents equality between measures. Solid grey line represents the regression line. Shaded grey area represents 95% confidence interval around the regression line.

Importantly, the LDA performed very well when considering mouse-level performance indicators. The Ad.ratio represents the ratio of the LDA-derived Ad count over the CALL-provided Ad-count; the nAd.ratio is a similar indicator for nAd features. If the LDA was, in practice, perfect, these ratios would be of exactly 1 (although it should be noted that the converse is not true, and a ratio of 1 does not correspond to perfect performance). We observed that the majority (the “most average 50%”, as indicated by the gray boxes in **Fig. 5B**) of validation dataset Ad.ratios were between 0.919 and 1.051, with a median of 0.984, while the whole dataset achieved an Ad.ratio of 0.941. The nAd.ratio performed arguably even better, with the majority of validation dataset nAd.ratios being between 0.965 and 1.054 with a median of 1.010, compared to an overall dataset performance of 1.04. Full indicator quantiles are given in **Tables S3 & S4**, with the 0% and 100% quantiles indicating the minimum and maximum values, that is, 0% and 100% of datasets below the indicated values, respectively. Taken together, these results indicate that despite the presence of a low frequency of inaccurate tumour callings, the estimated mouse-level tumour count is highly accurate.

Both the adenoma numbers and the total adenoma areas calculated by LDA showed high correlation to the values obtained using DRAW or CALL. Concordance between LDA and CALL was very good, in general with LDA obtaining only slightly less Ad counts than CALL, as shown by the regression line & confidence region thereof in **Fig. 5C**. Quite interestingly, the total Ad area was a much more accurate and consistent mouse-level measure compared to the number of Ad, as evidenced by the tight regression line in **Fig. 5E**. Unsurprisingly, the LDA approach yields mouse-level measures closer to those of CALL rather than that of DRAW, as it was trained and used on adenoma callings from the CALL approach (**Fig. 5D for counts and 5F for area**); however, all three approaches generate similar tumour number and total tumour area measures, indicating a good predictive value across the three methods.

### Adenoma area is a valuable measure of tumour burden

Many previous papers have used total tumour number as the only measurement of tumour burden to assess the effects of various treatments on *Apc^Min^* mice (for example,^13–15^). However, this does not take into account the size of the tumours, which can also be highly variable.

The automated method described here greatly facilitates the measurement of total adenoma area. We investigated how appropriate total adenoma area is as a measure of tumour burden. **Figure 6A** illustrates why total area should be measured and recorded: it presents two samples with identical tumour counts, but largely different tumour sizes. Biologically, larger tumours in the colon have been shown to be associated with shorter patient survival, showing the importance of considering tumour size as well as number in response to treatments^16,17^ Furthermore, **Fig. 6B** illustrates that, in a sample of 63 mice evaluated using the DRAW method, the average area of each tumour varied between different sections of the intestinal tract, with tumours in the LI being significantly larger on average than SI tumours. We also examined the correlation between total adenoma area and adenoma count in the SI. As shown in **Fig. 6C**, the correlation between adenoma area and count was high, but the spread increased with tumour number, thus reinforcing the utility of both measurements in evaluating tumour status. Finally, we correlated the number and total area of tumours in the SI to spleen weight, which represents a good surrogate measure of health status in *Apc^Min^* mice. Total tumour area in the SI was a better correlate of spleen weight than tumour number (**Fig. 6D**), even when excluding a potential outlier (R^2^= 0.36 vs. 0.43). We argue that these observations, taken together, demonstrate the need to evaluate tumour area in addition to tumour count.

**Figure 6.**
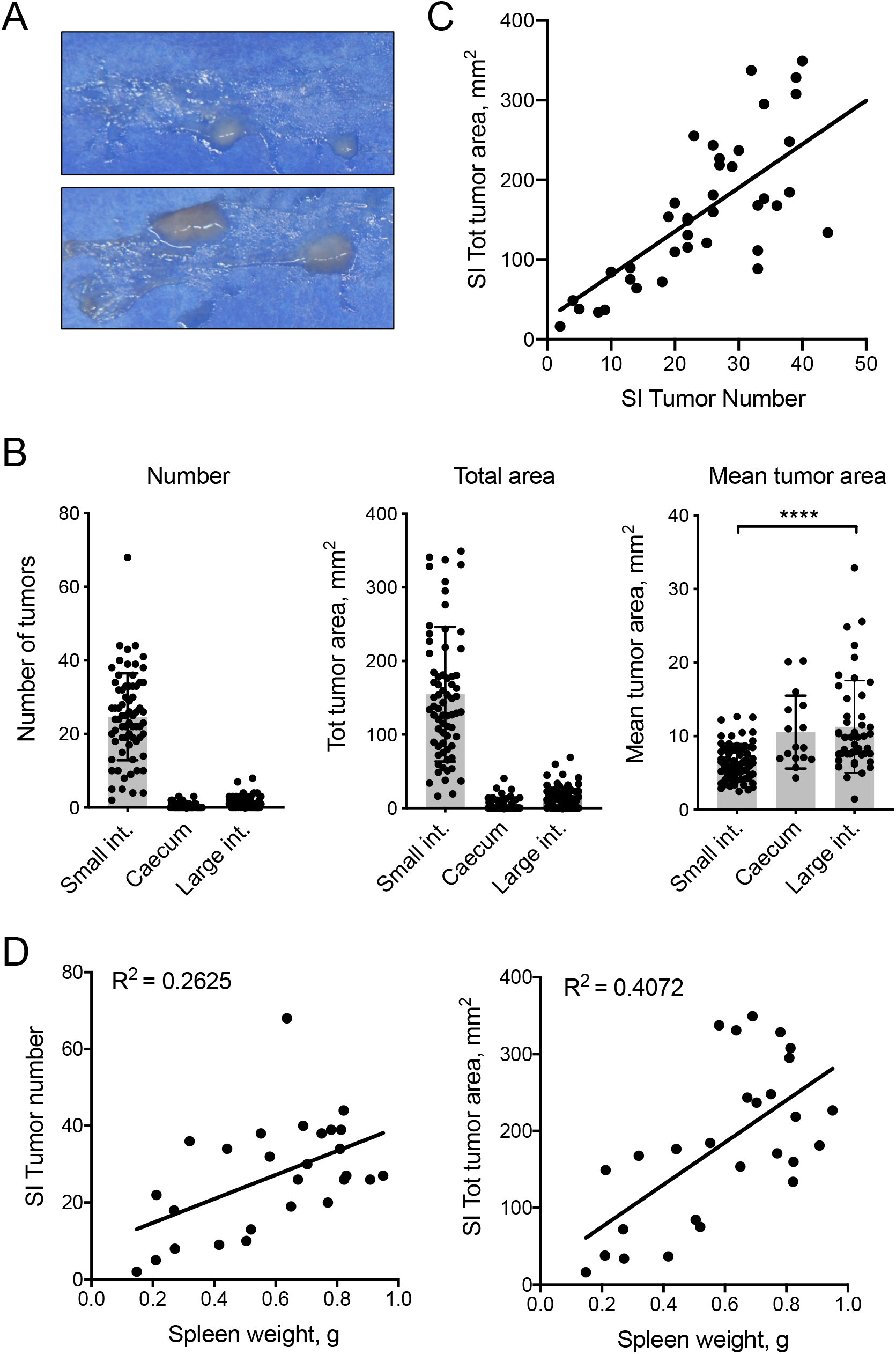
Total tumor area is an informative measure of tumour burden in Apc^Min^ mice. (**A**) Duodenal samples from two Apc^Min^ mice, each with two tumours. Note the large difference in tumour sizes between the two samples. (**B**) Bar graphs show the mean Number, Total area and Mean tumor area of tumours in different locations of the intestinal tract, +/− SD. Tumors were identified and measured using the DRAW method in a sample of 70 Apc^Min^ mice. Each dot represents a single mouse. ****: p<0.0001 as determined using a Kruskal-Wallis test with Dunn’s multiple comparison test. (**C**) Correlation between tumour count and size in LDA-called features in the SI of 35 mice. The dotted line represents the regression line. (**D**) Linear regression analysis of spleen weight vs. SI tumor number (left panel) or total area (right panel) in the SI of 27 mice for which spleen weight was available. Each dot represents one mouse. Data are from Figure 4.

### Utility of the total adenoma area measurements as assessed by LDA

To evaluate the usefulness and comparability of the tumour burden measures established by the DRAW and LDA approaches, we compared their power to discriminate between tumour burdens in mice of different ages (147 days or younger versus older than 147 days at the time of sacrifice, which are expected to have different tumour burdens) as a proof of principle. These comparisons are illustrated in **Fig. 7A-D**. Younger mice show a significantly lower number of Ad and total Ad area than older mice, in both the DRAW and LDA method, thus validating that both manual and automatic classification of SI features can distinguish between lower numbers and area of adenomas. Unsurprisingly, differences were much more pronounced for the area measures than the counts, further illustrating the utility of area as a measure of tumour load.

**Figure 7.**
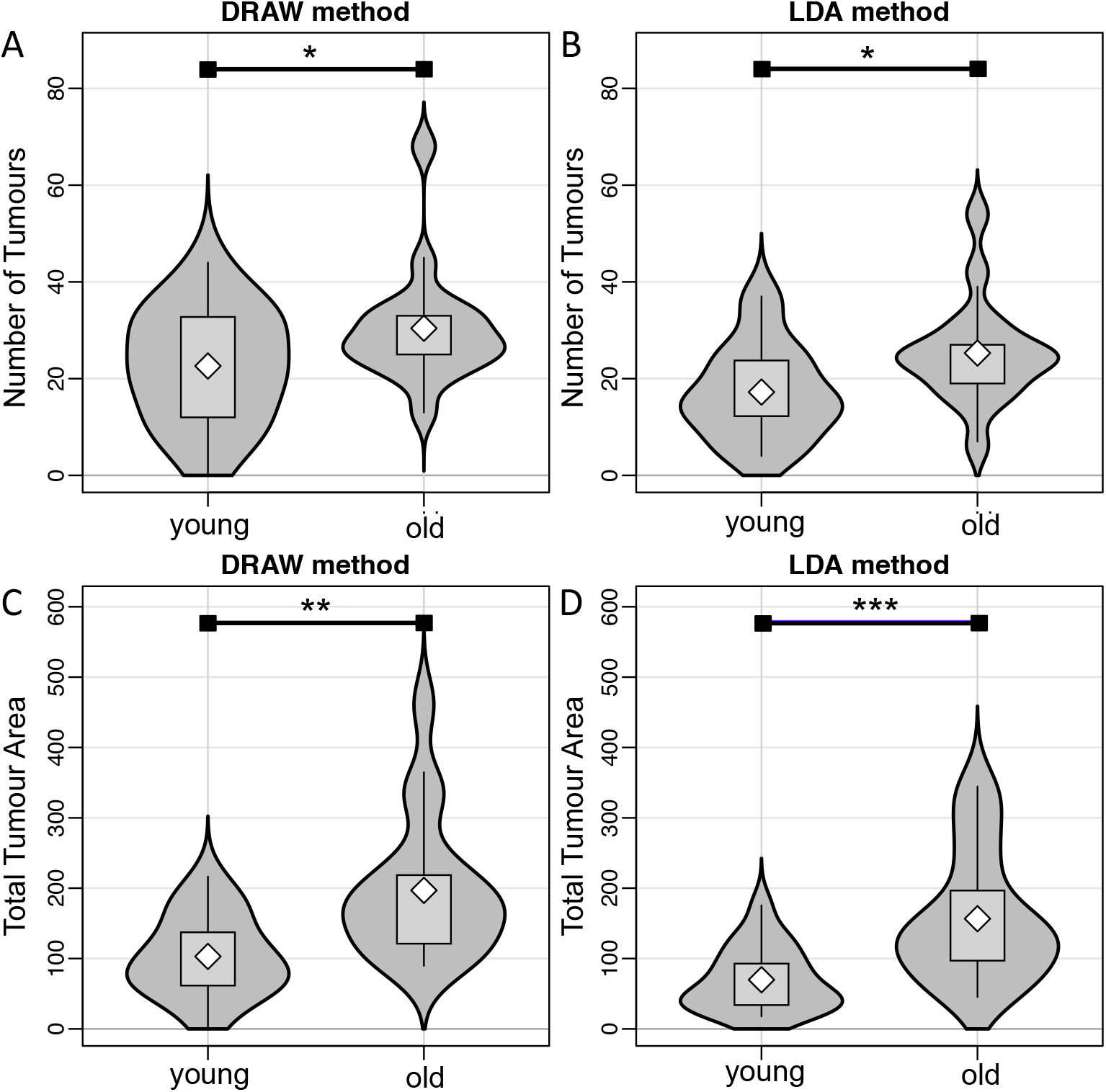
The DRAW and LDA methods both differentiate tumour number and total tumour area in young vs. old mice. 35 *Apc^Min^* mice were sacrificed when anaemic and then split by age: ‘Young’ (n=18) range from 1-147 days, while ‘Old’ (n=17) range from 147-214 days. (**A, B**): Violin plots of the number of tumours enumerated by the DRAW method (**A**) and the LDA method (**B**). (**C, D**): Violin plots of total area of tumours calculated by the DRAW method (**C**) and the LDA method (**D**). Stars indicate significance at the 5% level for approximate one-tailed Mann-Whitney-Wilcoxon tests (*: p<0.05, **: p<0.01, ***= p<0.001).

## DISCUSSION

We have developed a standardized protocol for first preparing and photographing mouse SI samples, then for the manual (using the DRAW approach) or automatic (using an ImageJ macro, *FeatureCounter)* identification of interesting image features (CALL approach), and finally an LDA-based method for the automatic classification of said features as true Adenomas or not Adenomas. Taken as a whole, these strategies allow for the consistent, rapid and robust derivation of mouse-level tumour burden measures (both as adenoma count and total adenoma area) for subsequent analysis.

Each of the steps in this standardized protocol works towards reducing technical, mouse-, experimenter- and even institute-level bias and variability, thus increasing result comparability and reproducibility. Additionally, the benefits are synergistic: as already pointed out, more controlled sample preparation allows for more consistent feature identification; and more consistently-defined features make feature classification easier. To note, best results for training the LDA classifier would be expected by using training sets called manually by either a single experimenter (as in this study), allowing the LDA to “learn” the same cues as that experimenter, or by as many different experimenters as possible (preferably across the same mice) allowing the LDA to “learn” the common cues to all. Even with the best practice, however, the correct classification of image features by our LDA step was not perfect. Most certainly, each step of our proposed method can be further improved in future research. The use of diffuse lighting (such as a photography tent) at the photography stage would minimise reflections that can be picked up as image features. The *FeatureCounter* may be adjusted to detect less features in tumour-less images (for example, by increasing the threshold size to ignore small features), while the automatic classification may be adjusted or replaced with another machine learning methodology. For example, a GLMNET algorithm^18^ would allow the simultaneous selection and estimation of input variable coefficients, at the very least leading to more consistent, if not more accurate, results. More advanced machine learning algorithms, such as neural networks, are now being used in the analysis of images from pathological samples, with new quantification approaches becoming available (reviewed in^10^). In some cases, deep neural networks have been shown to deliver classifications that are as accurate as those of a specialist, as in the case of skin lesions^11^. Therefore, neural networks, of which LDA is a simple, single node example, have the potential to provide better classification of images such as those generated in this study. In any case, manual verification by an experienced researcher can be rapidly and easily associated with any of the protocols described here, and would be most conveniently carried out after LDA corrects the most evident misclassifications, such as those resulting from imperfect sample preparation or photography– although these are relatively rare once the technique is learned (3/120 in this study).

Regardless, our semi-automated strategy is faster, more reliable and also more flexible than previously used methods. Samples can be processed and analysed while fresh, or can be frozen and analysed later at a convenient time. Through the sample freezing step, “break points” are introduced into the experimental workflow, *i.e.* points at which the experimentation for a single sample can be suspended temporarily, while in traditional methods each sample is often prepared and counted the same day. The reduced time cost in tumour quantification can be another major benefit in the DRAW, CALL and LDA approaches. It is thus immediately apparent that, beyond the added flexibility, our automated strategy may earn a considerable sample preparation and counting time gain when many mice – especially heavily tumour-burdened ones – are being assessed. Furthermore, the preparation techniques are accelerated further when processing multiple samples at a time. Additionally, the wealth of data is higher using these approaches compared to the TRAD count method, where just tumour number, or cumulative tumour area in a small section of the intestine, is assessed. We also note that once digitized, the photographic information can be stored almost indefinitely, allowing the data to be revisited if need be, for example, after a *FeatureCounter* update, or after the implementation of a new classification methodology, or for meta-analysis. Finally, if the effort of generating a large LDA training set was not justified, the CALL and DRAW methods can be rapidly implemented, and are still quicker, more reliable, and producing more detailed data than the traditional method.

Several previous papers (for example,^13–15^) have only reported on total adenoma number, using this as the lone tumour burden measure to assess the effects of various treatments on *Apc^Min^* mice. However, this does not take into account the size or aspect of the tumours, which can be highly variable. For our part, we believe that adenoma count certainly cannot be used alone, as area can differ for identical adenoma counts, and its distribution changes between different segments of the mouse intestinal tract. The reasons for these similarities and differences are multiple. For example, early studies of the *Apc^Min^* mouse strain reported that adenomas develop mostly during early life and up to puberty, and their numbers did not increase after 100 days of age^19^. After this stabilisation in numbers, the adenomas have been observed to instead grow in size^20^, thus increasing tumour burden in a way not captured by adenoma count alone. Additionally, significant size differences have been found in some cases, demonstrating that area measures can provide additional information about treatments or exacerbating conditions^21^. For example, therapies may be effective at controlling adenoma growth without fully eradicating tumours, an effect that would be detected as decreased burden with little or no change in tumour number. We thus conclude that adenoma area, and potentially other measures, are of sufficient importance and value to warrant the use of new methods to facilitate collection of such information. As adenoma number is still generated using our approach, comparisons to previous studies remain possible. Of note, with our ImageJ feature-based approach, it is possible to derive several aggregate measures (for example, average adenoma greyscale value per mouse, as listed in parameters in **Supplementary Table 2**) that might relate back to tumour burden or other biological indicators of interest. Further research in this direction may yield interesting insights.

In conclusion, we propose a semi-automated method to rapidly quantify tumour number and associated tumour burden measures that will help alleviate biases, along with reproducibility and consistency problems, which currently hamper efforts to interpret results across the *Apc^Min^* mouse literature. Our method is convenient, can be adapted to provide measurements of several tumour characteristics, and will facilitate the use of *Apc^Min^* mouse intestinal adenoma model in a variety of applications.

## METHODS

### Animals

C57BL/6J-*Apc^Min^ (Apc^Min^)* mice were purchased from The Jackson Laboratory (Bar Harbor, ME) and bred in SPF conditions at the Malaghan Institute of Medical Research by mating C57BL/6J-*Apc^Min/+^* males with Wild-type (WT) C57BL/6J *(Apc^+/+^)* females. *Apc^Min/+^* and WT offspring were identified by PCR and were both used in experimental conditions and pipeline development. Water and standard laboratory chow were available *ad libitum.* All mice were checked regularly for signs of anaemia and sickness, and were euthanized for tissue collection if they developed pallor, low haematocrit (< 20%), weight loss, slow movement and/or hunched posture.

All experimental protocols were approved by the Victoria University of Wellington Animal Ethics Committee, and were carried out in accordance with the Victoria University of Wellington Code of Ethical Conduct.

### Tissue preparation

Mice were euthanized and the entire intestinal tract was extracted and sectioned into the SI, caecum and LI. Special care was taken to remove as much mesenteric fat as possible. Sections were washed thoroughly using PBS, drained, and analysed immediately or frozen in 6 well plates at −80°C until further use.

### Photography

For image analysis, SI tract sections were thawed (if frozen) and spread out in a thin horseshoe shape on pieces of Steel Blue Germination paper (Anchor Paper Company, St Pauls, MN, USA) approximately 25×10 cm in size. This colour was selected to enhance the contrast between adenomas and the rest of the intestine. Once laid out on the paper, the SI was cut into 2 equal pieces. Each piece was then cut longitudinally along the tube, opened and edges spread flat using the edge of curved tweezers. Mucus and intestinal contents were removed by spraying PBS on the tissue preparation, revealing any adenomas present. The preparation was then photographed with a Panasonic Lumix G Vario DMX-G5W and a 45-150 mm lens with additional 4x filter (Marumi, Japan), with a white ruler in shot. Multiple pictures were taken and stitched together to reconstruct an image of the entire SI using the software *Hugin* 2013.0.0^22^.

For the LI and caecum, a similar strategy was undertaken, where the tissue was placed on the same type of Steel Blue Germination Paper as the SI, cut longitudinally (with multiple cuts needed for the caecum), spread as flat as possible with special care taken to flatten tissue near an adenoma in the caecum, and photographed with the white ruler in shot. Both the LI and the caecum are small enough that they could be captured in one photograph.

### Manual delineation of tumours in images (DRAW approach)

In order to enumerate and measure the area of tumours in the stitched images of SI, LI and caecum, we used the Java-based image processing programme “ImageJ” (https://imagej.net,^12^), which is freely available and able to analyse images in a variety of formats. Full detail on tissue preparation, photography and analysis can be found at https://gitlab.com/gringer/featurecounter/blob/master/Sample_Photography.pdf.

Images were scaled using a small macro and the white ruler in shot as a reference. ImageJ’s ‘freehand selection’ function was then used to manually delineate visually-identified image regions corresponding to adenomas. A scaled mask image was created using ImageJ’s ‘create mask’ function, and was analysed with the ‘analyze particles’ function to generate adenoma numbers and measurements such as area. This is referred to as the “DRAW” approach.

### *FeatureCounter*, an ImageJ macro for the automatic identification of image features

In order to automate the identification of image regions potentially corresponding to adenomas from the photographs of intestinal sections as described in the DRAW approach, we developed a more extensive ImageJ macro, called “*FeatureCounter*”, focusing on SI sections as these contain the large majority of the tumours that develop in *Apc^Min^* mice. First, *FeatureCounter* subtracts the blue background, leaving a grey scale image. It subsequently performs automatic thresholding, before despeckling the image according to the parameters listed in **Table S1**. This leaves areas of over 0.2mmsq in size, or “image features” that are potentially tumours. The “analyse particles” function within ImageJ measures 22 variables for each feature: Area, Perimeter, Mean, StdDev, Mode, Min, Max, Median, Skew, Kurt, Major, Minor, Angle, Circularity, AR, Round, Solidity, Feret, FeretAngle, MinFeret, IntDen, and RawIntDen. The details of these measures and their processing can be found in **Table S2.** *FeatureCounter* was optimised to work on the SI due to its smooth and regular surface. It does not perform as well at quantifying tumours in the LI, where the surface of the intestinal wall is ridged, or in the caecum, where the tissue does not spread out flat particularly well. As the number of tumours in the caecum and LI rarely exceeds 3 (mean and SD of LI and caecum is 1.81 ± 2.00 and 0.41 ± 0.75 respectively), these tumours can be quickly and accurately quantified manually from photos using the DRAW approach. Therefore, further work to optimize *FeatureCounter* performance on the LI and caecum did not seem warranted, and was not pursued.

### Manual validation of tumour features (CALL approach)

Image features identified by *FeatureCounter* can be manually validated. After running the macro, a user can manually assign or “call” which features are tumours, referring to them as “Adenoma” (Ad) or “not-an-Adenoma” (nAd) or, for unclear features, ‘Not Assigned’ (NA). In our study, there were relatively few NA features, and they were consequently excluded from further analyses. We refer to this approach as the “CALL” approach.

Of further interest, the image feature measures obtained from *FeatureCounter* can be leveraged in a machine learning algorithm to automatically determine which features are tumours and which are false positives. Such a machine learning algorithm would require a gold-standard “training dataset”, i.e., a dataset of image features, their measurements, and a prior validation of which features are indeed tumours or not, to learn tumour-specific patterns. The CALL approach can be used to generate such a training set.

### Linear Discriminant Analysis (LDA) for automatic classification of image features

Using the image feature measurements from *FeatureCounter* and a training dataset as prepared using the CALL approach above, a machine learning technique can be used to attempt to automatically separate tumour features from non-tumour features using the feature measurements. LDA is one such supervised classification technique. It determines discriminant functions – or the optimal linear combinations of the various input variables (here: the 22 feature measures) – that can be used to classify statistical observations (here: image features) into different classes (here: Ad or nAd). In our implementation, the *squares* of the input variables were included as further input variables, as this allows quadratic separations within the original variable space. All data were analysed within the R statistical programming framework^23^.

LDA is sensitive to several influences, including 1) extreme non-normality in input variable distributions and 2) extreme outliers in input variables. For these reasons, it is recommended to pre-process the input variables. We manually examined the distributions of the feature measure variables per class, and applied log10 transformations, shifted log10 transformations, and imposed certain filters, as described in **Table S2**.

The applicability of LDA to the transformed feature data was first evaluated by performing a Principal Components Analysis (PCA) with package *FactoMineR*^24^, the assumption being that if the major axes of variability in the measurement data cannot segregate the classes even partially, there is no point in performing an LDA and more advanced machine learning techniques need to be used. The LDA was then performed using the *lda* function in the R package *MASS*^25^ for features with no missing values. A link to the R script used to run the LDA can be found in the Supplementary materials. We then proceeded to investigate the performance of our LDA at two levels, described below: at the feature level (checking whether the classification performed well) and at the mouse level (checking whether, in practice, the methodology allowed for accurate tumour counting and area quantification).

### LDA feature-level and dataset-level performance

We compared the LDA’s feature-level predictions to the adenomas selected using the CALL method, which were considered “true” adenomas in this instance. We considered as indicators of the LDA’s performance the True Positive Rate (TPR, or Sensitivity, here defined as the proportion of all true Ad that were also identified as adenomas using LDA), the Positive Predictive Value (PPV, or the proportion of the LDA-identified adenomas that were indeed Ad), and the Accuracy (the proportion of all features correctly identified as Ad or nAd). Similar calculations were done for the nAd classes.

As indicators of dataset-level performance of the CALL and LDA adenoma callings, we counted the number of Ad and nAd calls, and calculated the ratios of the number of LDA-predicted Ad and nAd over the number of CALL-provided Ad and nAd (Ad.ratio and nAd.ratio, respectively). An LDA with perfect performance would generate ratios of exactly 1, although a value of 1 is not necessarily indicative of perfect performance.

### LDA validation

To assess the robustness of the LDA’s results, we performed a large validation experiment with a complex re-sampling scheme inspired by those of mixed modelling/multi-level models. We chose to randomly sample mice (with replacements, *i.e.* a same mouse can be sampled more than once) from the 117 with appropriate data, including all their image features in each validation dataset. Mice continued to be sampled until a) at least 12 mice (about 10.3% of the total) had been sampled, and until b) at least 750 features (23.5% of total) had been sampled. Indeed, as the choice of the feature number parameter in the re-sampling scheme strongly influences the performance indicators, we empirically determined that a minimal feature count of 750 presented the best trade-off between sample size and indicator performance (**Supplementary Fig. 4**). Additionally, to ensure some measure of class balance, only datasets with a composition containing at least 30% Ad features and 30% nAd features were retained. A total of 4000 validation datasets were generated (computationally representing the equivalent number of ‘experiments’ of normal *Apc^Min^* and WT animals), and each was used to train a separate LDA. For each validation LDA model, feature-level performance indicators (Accuracy, TPR, PPV) and dataset-level performance indicators (Ad.ratio and nAd.ratio) described above were derived using the whole dataset. For all indicators, we established their quantiles of interest (0, 5, 25, 50, 75, 95, 100%) to compare to the values obtained on the full dataset LDA.

### Statistics used to compare mouse-level results

Comparisons of mouse data (weight, tumor numbers etc) used the Mann-Whitney U test or a Kruskal-Wallis test followed by Dunn’s multiple comparison test, and were performed using Prism 8.0 GraphPad software.

To compare adenoma results at the mouse level (counts, total areas) obtained using different methods (CALL and LDA), we used Deming regression, a statistical technique used for comparing two measurement methods for a same quantity, where *both* measurements are assumed to have measurement error (typical linear regression only assumes error in the outcome variable). We used the *mcreg* function implemented in package *mcr*^26^ assuming a variance ratio of 1, and using bootstrapping (n=999, ‘Bias-corrected and accelerated’ method) to obtain a regression curve confidence area.

### Availability of Data and Materials

The *FeatureCounter* ImageJ macro is freely available to download from https://gitlab.com/gringer/featurecounter/together with instructions for photography, and macro installation, some examples of tumour images, and the R code for running the LDA. The datasets generated during the current study are available from the corresponding author on reasonable request. Tumour images are available from Zenodo repository. doi:10.5281/zenodo.3365777.

## Acknowledgements

We sincerely thank all staff at the Malaghan Institute of Medical Research Biomedical Research Unit for maintenance of the *Apc^Min^* mouse strain. We thank Professor Terry Speed, (Walter and Eliza Hall Institute of Medical Research) for helpful discussions about statistical analysis. We also thank the Free and Open Source Software community for access to a number of programs used over the course of this project. This work was funded by the A.M. Duncan Bequest to the Malaghan Institute of Medical Research, and funding from the NZ Cancer Society and the Health Research Council of NZ to FR.

## Author Contributions

ALS developed and performed gut preparations, counted adenomas manually and classified the automatically identified features, and wrote the manuscript. AATS carried out statistical analyses of image features. KAW carried out adenoma validation and generated adenoma data. SK and JY carried out TRAD quantifications. DAE developed the *FeatureCounter* macro. FR supervised the project, provided support and suggestions for investigations, and edited the manuscript. All authors provided suggestions and approved the final manuscript.

