## Supplementary Figures and Tables for "A semi-automated technique for adenoma quantification in the *Apc^Min^* mouse using *FeatureCounter*"

**Supplementary Information**

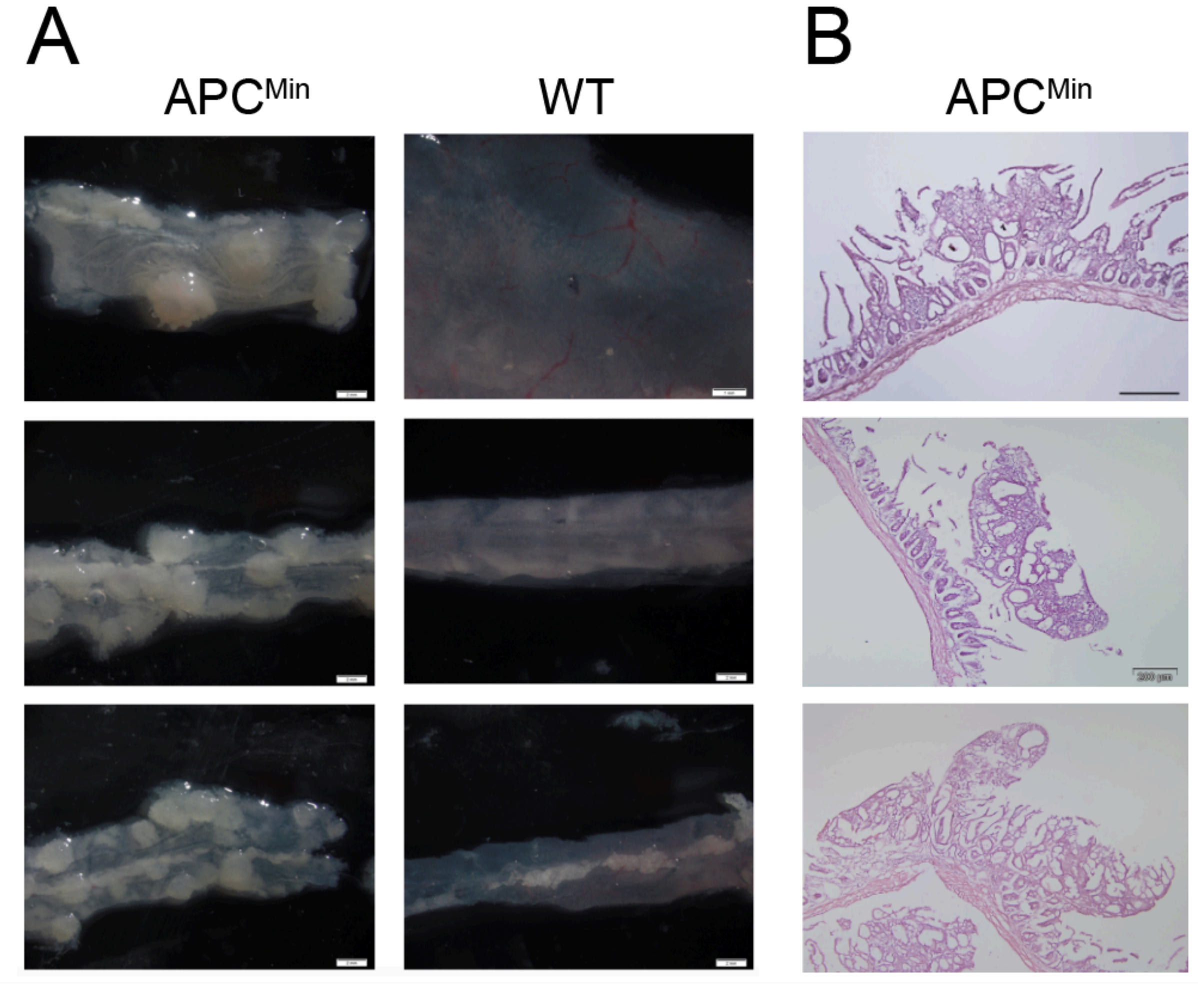


**Supplementary Figure 1. Intestinal tumors in APC/Min mice and WT littermates.** APC/Min mice were sacrificed at the first sign of anemia; WT littermates were sacrificed at the same time. Intestinal tissue was collected for processing and tumor visualization as indicated.

(**A**) Intestinal tissue was opened longitudinally, washed with PBS, then immediately examined at low magnification under a stereomicroscope. Images show Duodenum (upper row), Upper jejunum (middle row) and Ileum (lower row). Size bars correspond to 1 mm (WT duodenum) or 2 mm (all other images).

(**B**) Intestinal tissue was collected and examined macroscopically. Areas corresponding to putative tumours were removed, fixed in formalin, paraffin embedded, sectioned and stained with H&E, and examined microscopically. Size bars correspond to 200um.

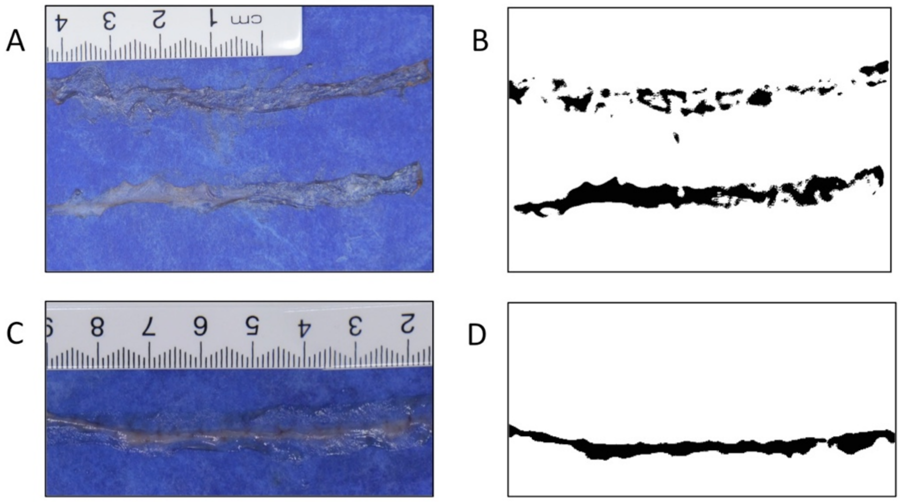


**Supplementary Figure 2. The *FeatureCounter* macro can be misled by low quality sample preparations.**

Intestinal tissue from APC+/+ (WT) mice was collected and processed as described in the methods

(**A**) Photograph of curled edges (top section) and dried (bottom section) SI preparations. (**B**) *FeatureCounter* generated tumour masks identify curled edges (top section) and dried areas of SI as interesting features that might be Ad. (**C**) A section of SI with remaining excess fat. (**D**) Corresponding tumour mask showing that excess fat is identified as an interesting feature by *FeatureCounter*.

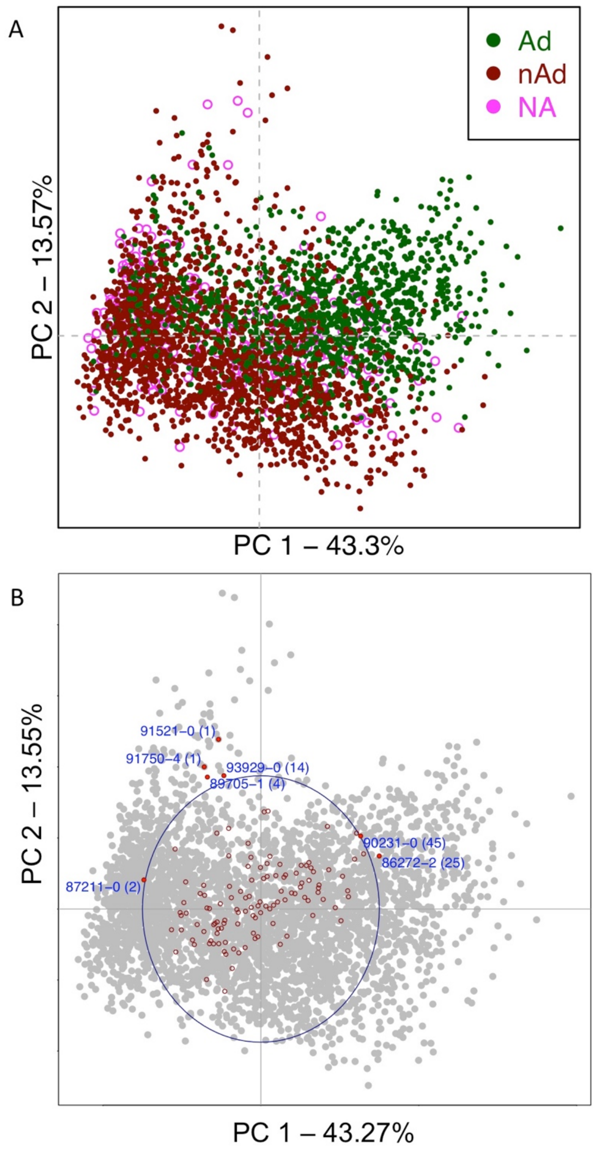


**Supplementary Figure 3. Principal Components Analyses of LDA-generated image features**

PCA was applied to features with 22 measures (after transformation and filtering) from 120 mice. (A) PCA applied to 3188 Adenoma (Green) and Non-Adenoma (Red) features. Additionally, 259 Not Assigned (Pink) features are illustratively projected onto the plot. (B) PCA applied to 3447 (Adenoma, Non-Adenoma and Not Assigned) features from 120 mice. Features are indicated as grey dots. Mouse barycentres are indicated as red circles. Red dots with blue labels indicate mice with distinctly non-central barycentres, i.e. potentially biased mice. Parentheses give feature counts in those mice.

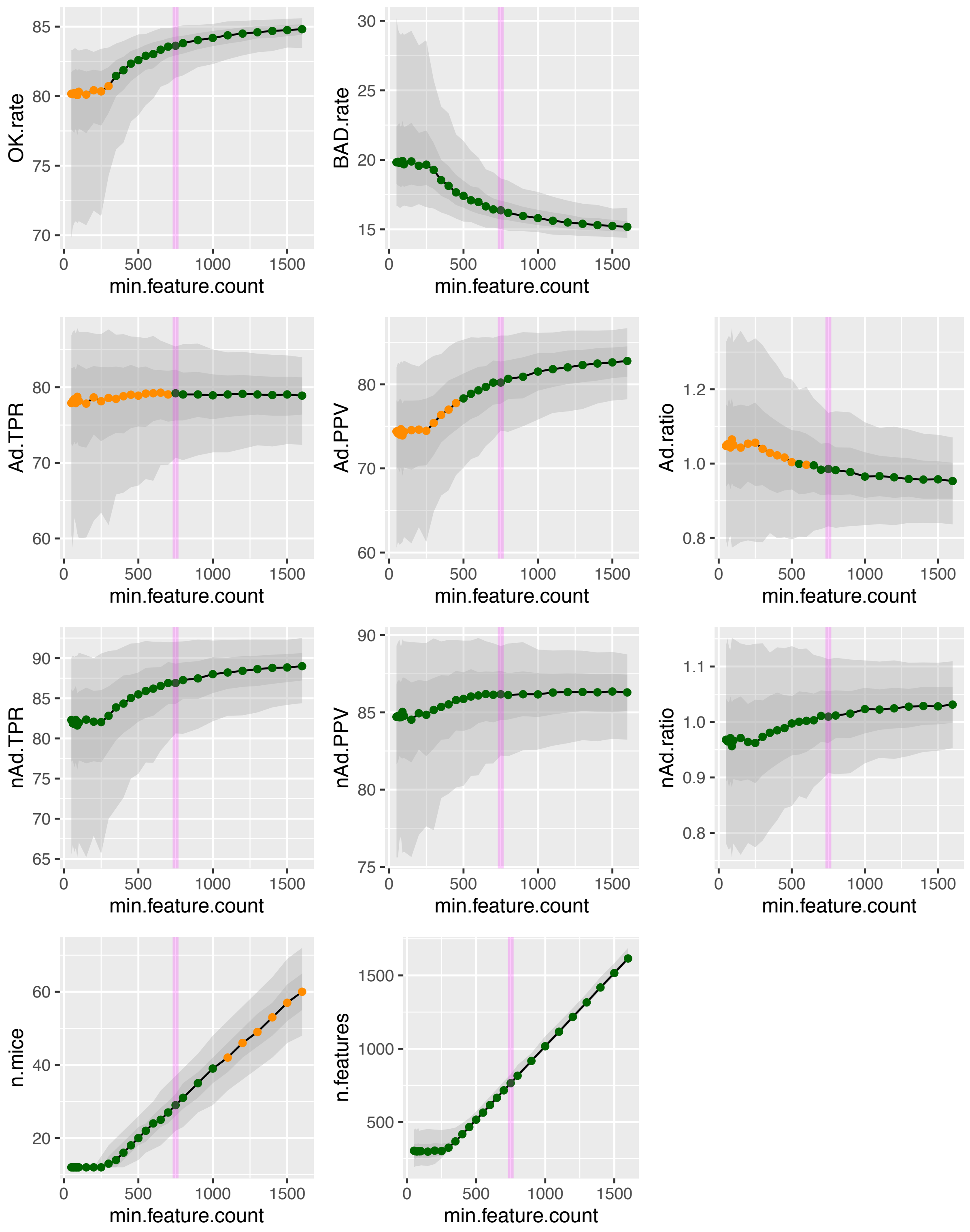


**Supplementary Figure 4: Selecting a minimum feature count for bootstrapping resampling training datasets.**

For several values of minimal feature count (broadly taken between 50 and 1600), performance metrics were calculated across 1000 validation datasets, built as described in the methods. Each plot represents the 5% - 95% quantiles (light grey area), 25%-75% quantiles (dark grey area), and 50% quantile (i.e. median, black line) for one metric. The dots along the line are coloured according to selection conditions we established (OK.rate 5$^{th}$ percentile ≥ 75% ; Ad.TPR 5$^{th}$ percentile ≥ 70% ; Ad.PPV 5$^{th}$ percentile ≥ 70% ; Ad.ratio between 0.80 and 1.20 ; n.mice 95$^{th}$ percentile ≤ 50) to help guide our selection of a minimal feature count (green: passes condition; orange: fails condition) necessary for appropriate bootstrapping resampling-based validation. The vertical magenta line indicates our final choice for minimum feature count to use in the validation sets. OK.rate: percentage of correct predictions over total predictions; BAD.rate: complement to OK.rate; Ad.TPR: True Positive Rate on Adenoma predictions; Ad.PPV: Positive Predictive Value on Adenoma predictions; Ad.ratio: ratio of number of predicted Adenomas over actual number of Adenomas; nAd.TPR, nAd.PP, nAd.ratio: same as for Adenomas, applied to not-an-Adenoma features; n.mice, n.features: number of mice & number of total features included in validation dataset

**Supplementary Table 1:** Automatic thresholding and filtering to select "image features" that are potentially adenomas.

| Step | Action |
|---|---|
| 1 | Min/max range set to 60-200 to exclude background noise |
| 2 | Threshold set to 75 (of 255 grey levels) |
| 3 | Remove dark outliers of radius 20 with threshold 50 |
| 4 | Remove light outliers of radius 20 with threshold 50 |
| 5 | Dilate to increase edge diameters slightly |
| 6 | Watershed to split multiple joined features into component parts |
| 7 | Analyze Particles with "size=0.2-Infinity" |

**Supplementary Table 2:** ImageJ “analyse particles” variables measured by *FeatureCounter* and their processing

| Measurement | Description | Shift? | Log-transform? | Min. | Max. |
|---|---|---|---|---|---|
| Area | Area of feature | | Yes | | |
| Perimeter | Perimeter of feature | | Yes | 0 | 2 |
| Mean | Mean grey value within feature | | | | |
| StdDev | Standard deviation of grey value within feature | | | | |
| Mode | Modal grey value within feature | | | | |
| Min | Minimum grey value within feature | Yes | Yes | | |
| Max | Maximum grey value within feature | | | | |
| Median | Median grey value within feature | | | | |
| Skew | 3rd order moment about the mean of grey values in the feature | Yes | Yes | 0.25 | 1 |
| Kurt | 4th order moment about the mean of grey values in the feature | Yes | Yes | 0 | 1.5 |
| Major | Major axis of an ellipse fit to the feature | | Yes | | |
| Minor | Minor axis of an ellipse fit to the feature | | Yes | | |
| Angle | Angle between the axes of the ellipse fit to the feature | | | | |
| Circularity | Measurement of circularity of the feature | | | | |
| AR | Aspect ratio of the ellipse fit to the feature - major over minor | | Yes | | |
| Round | Roundness - inverse of aspect ratio | | | | |
| Solidity | Ratio of area to area of a convex hull fit to the feature | | | | |
| Feret | Segment of longest distance between two points along feature boundary | | Yes | -0.3 | 1.5 |
| FeretAngle | Angle of the Feret segment | | | | |
| MinFeret | Minimum caliper diameter of the Feret segment | | Yes | -0.5 | 1.5 |
| IntDen | Integrated density: estimated sum of grey values in the feature | | Yes | | |
| RawIntDen | Raw Integrated density: sum of grey values in the feature | | Yes | | |

Feature measurements returned by ImageJ's "Analyse Particles" function within *FeatureCounter*, with subsequent transformations and filtering information for use. C.F. https://imagej.nih.gov/ij/docs/guide/146-30.html (link pulled 12/02/2019) in the Linear Discriminant Analysis.

**Supplementary Table 3.** LDA indicator quantiles for Adenomas across bootstrapping datasets

| Quantile | Ad.TPR | Ad.PPV | Ad.ratio | Mice | Features |
|---|---|---|---|---|---|
| 0% | 57.62 | 65.78 | 0.68 | 15 | 750 |
| 5% | 70.13 | 73.95 | 0.82 | 22 | 751 |
| 25% | 75.82 | 77.84 | 0.92 | 26 | 757 |
| 50% | 79.12 | 80.25 | 0.98 | 29 | 766 |
| 75% | 82.17 | 82.71 | 1.05 | 33 | 781 |
| 95% | 85.77 | 85.76 | 1.15 | 37 | 827 |
| 100% | 91.01 | 90.64 | 1.36 | 45 | 959 |

Quantile: Target quantile (in %) across all 4000 cross-validation datasets comparing LDA-identified adenomas to the CALL Adenomas. Ad.TPR: True Positive Rate of Adenoma callings; Ad.PPV: Positive Predictive Value of Adenoma callings; Ad.ratio: Ratio of the number of Adenomas called by LDA over the number of True (CALL) Adenomas; Mice, Features: Number of mice and features sampled in the cross-validation set (with replacement), respectively.

**Supplementary Table 4.** LDA indicator quantiles for not-Adenomas across bootstrapping datasets

| Quantile | nAd.TPR | nAd.PPV | nAd.ratio | Mice | Features |
|---|---|---|---|---|---|
| 0% | 69.51 | 75.65 | 0.76 | 15 | 750 |
| 5% | 79.99 | 81.99 | 0.90 | 22 | 751 |
| 25% | 84.39 | 84.59 | 0.97 | 26 | 757 |
| 50% | 87.01 | 86.13 | 1.01 | 29 | 766 |
| 75% | 89.31 | 87.64 | 1.05 | 33 | 781 |
| 95% | 92.14 | 89.46 | 1.12 | 37 | 827 |
| 100% | 95.70 | 92.41 | 1.22 | 45 | 959 |

Quantile: Target quantile (in %) across all 4000 cross-validation datasets comparing LDA-identified not-Adenomas to the CALL not-adenomas; nAd.TPR: True Positive Rate of not-Adenoma callings; nAd.PPV: Positive Predictive Value of not- Adenoma callings; nAd.ratio: Ratio of the number of not-Adenomas called by LDA over the number of True Adenomas; Mice, Features: Number of mice and features sampled in the cross-validation set (with replacement), respectively.

List of materials available from GitLab (https://gitlab.com/gringer/featurecounter/):

| MD5 Hash | Path | Description |
|---|---|---|
| 89aaa67921d2a58181d74c492590c039 | ./Autoscale.ijm | Macro to add an image scale when it contains a horizontal ruler |
| 56aa13bdc1b4330debcc63a11a297d42 | ./MeasureFeatures.ijm | Macro to isolate, mark, and calculate statistics for interesting image features |
| 7ac61644fadd7cb6680e9e27c4b52c79 | ./SubtractBlue.ijm | Macro to subtract the blue background from an image |
| e3982ca783a403b770382b6c771cebcd | ./LICENSE.md | Software license (ISC / free software, non-copyleft) |
| acaae6bd455642f85e8b205578d2b6ff | ./README.md | "Feature Counter -- ImageJ macros to help identify tumour-like areas of an image" |
| bf5eede59380b4e7f1aef43e53509a4a | ./Sample_Photography.pdf | PDF for macro documentation |
| 0498dcadabca6e7906765752861258fc | ./Sample_Photography.tex | Text-file source for macro documentation |
| d536a191a38ac41c9989d85b1fc54dc5 | ./examples/images/SI81355-1.jpg | Example image: negative, light background |
| ad2368cc180edeed0627ac570fb2e929 | ./examples/images/SI90230-3.jpg | Example image: negative, medium background |
| 05da5b15e5b75397265728905861d37d | ./examples/images/SI90230-4.jpg | Example image: medium to high adenoma burden, mostly compound features of large size |
| 2c20781eb489003e7420ba1e40e75a89 | ./examples/images/SI90231-0.jpg | Example image: medium to high adenoma burden, mostly separated features of medium size |
| 8eba38b142e27d7f64983027f42c82c9 | ./examples/images/SI92332-0.jpg | Example image: low adenoma burden, blurry image, mostly separated features of medium size |
| a0ebdffcc17cd518c3e1bdbcce3af263 | ./examples/images/SI92335-0.jpg | Example image: negative, medium background, slanted ruler |
| a3e974619c14bfdeb315cc3fc26765a8 | ./examples/images/SI93929-3.jpg | Example image: negative, medium background, chopped off ruler at one side |
| dace1dcb7634798862f756a6fc17ff50 | ./examples/images/SI93930-1.jpg | Example image: negative, medium background, high light reflection |
| b08cbb0fdbabc0025aa122cf069499d3 | ./examples/images/SI94445-0.jpg | Example image: low adenoma burden, mostly compound features of large size |
| f9441940c36c28b8d8e55d54886465bf | ./examples/images/SI94445-1.jpg | Example image: low adenoma burden, lots of reflection and non-adenoma features |
| 34f298c079ff485840f926ad5efd573e | ./examples/images/SI94643-1.jpg | Example image: low adenoma burden, mostly separated features of large size |
| 545a7a3919399950dd5868ccde4e1670 | ./examples/images/SI95080-2.jpg | Example image: low adenoma burden, mostly separated features of large size |
| 14c46dbb85db82ab197dbc02ab48cd4a | ./examples/images/SI95779-2.jpg | Example image: low adenoma burden, mostly separated features of large size |
| 63bd6755a83698122130e547fddd6086 | ./examples/SimpleLDA.R | Example R code to carry out an LDA on feature statistics |
| 3fad2526eebbdb2d3f9a5c63800d043c | ./examples/FeatureData. | Example input for LDA code |
| 32a178fe39769c994e0fa796e26e676c | ./examples/lda_output.txt | Example output of LDA code |
| d1d1c5215c8341b77a45fcebf146fe18 | ./examples/Data_Apc_Min.xls | Example spreadsheet containing aggregate mouse-level metadata and image feature data |
| 021ebeeda7ceefdd5de284296679a8ba | ./illustrations/small_R-B_DkrPaint.jpg | Documentation illustration: after removing blue background |
| 8ba9d0139bb015e438d92f8c04e6b74b | ./illustrations/photography_setup.svg | Documentation illustration: camera setup for good image photography |
| 5fa337f208fbece1435ac204ceec1281 | ./illustrations/photography_setup.pdf | Documentation illustration: camera setup for good image photography (PDF version) |
| 89ef9ae1f234389a272d668f5eb2d513 | ./illustrations/stitching_setup.svg | Documentation illustration: image / photo layout for good stitching |
| c7927127856807b19088cb15c23147b6 | ./illustrations/stitching_setup.pdf | Documentation illustration: image / photo layout for good stitching (PDF version) |
| d8629e5eb538626b3041219d84d2ef4f | ./illustrations/small_foundRegions_DkrPaint.jpg | Documentation illustration: identified regions overlaid onto the original image |
| 57d06ec19f5b16bd69c2989242fdd493 | ./illustrations/small_DkrPaint.jpg | Documentation illustration: original image with highlighted ruler scale points |

Tumour images are available from Zenodo repository doi:10.5281/zenodo.3365777.
